# Hydrolysis of palm kernel meal fibre using a newly isolated *Bacillus subtilis* F6 with high mannanase activity

**DOI:** 10.1101/2024.06.19.599806

**Authors:** Wei Li Ong, Kam Lock Chan, Antonius Suwanto, Zhi Li, Kian-Hong Ng, Kang Zhou

## Abstract

High fibre content is the main limitation of using mannan-rich palm kernel meal (PKM) in feeding non-ruminant livestock. Microbial fermentation stands out as a cost-effective and environmentally friendly approach for hydrolysing fibre in lignocellulosic biomass. In this study, a *Bacillus subtilis* strain F6 with high mannanase secretion capability was isolated from an environmental source. Fermentation of PKM using strain F6 resulted in at least a 10% reduction in its neutral detergent fibre content. Notably, the strain exhibited a rapid response to PKM, with significant mannanase activity detected as early as 6 h, enabling fibre hydrolysis within a short fermentation period. Subsequent transcriptome analysis uncovered potential enzymes involved in PKM fibre degradation, and the purified recombinant enzymes were generated to assess their activity on PKM fibre degradation. β-mannanase GmuG demonstrated strong hydrolysis activity of PKM fibre, and its biochemical properties were determined. Overall, the study reported the isolation of a *B. subtilis* strain suitable for fibre hydrolysis of mannan-rich biomass, followed by an investigation to identify and characterize the enzyme responsible for fibre degradation.

**Graphical abstract:** 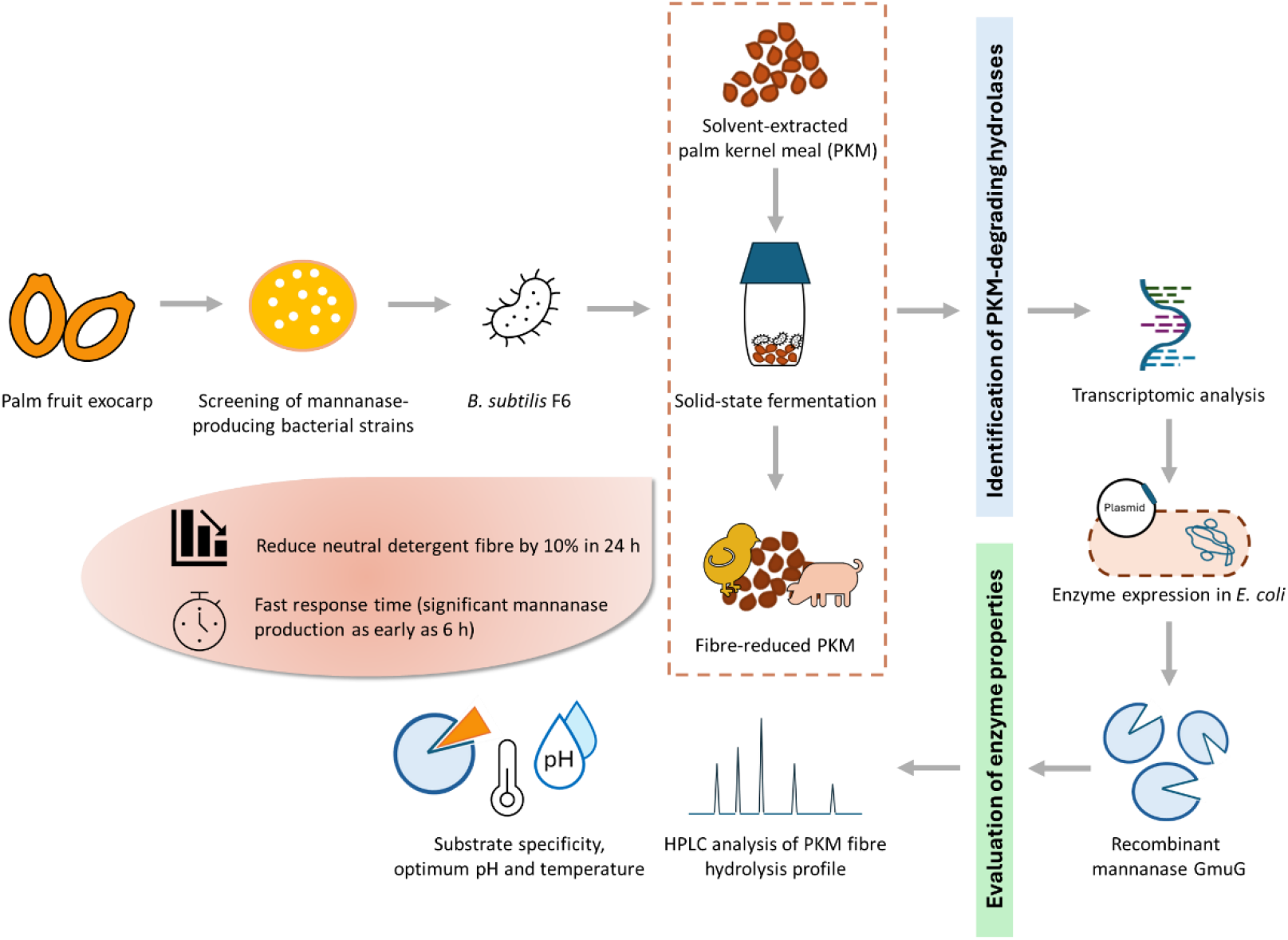

## 1. Introduction

Lignocellulosic biomass is a plant-derived material that primarily includes agricultural and forestry residues, energy crops, and yard trimmings (Xu & Li, 2017). This highly complex bioresource mainly comprises interwoven masses of cellulose, hemicelluloses, and lignin, with proteins, oils, and ash making up the remaining fraction (Sahay, 2022; Wei et al., 2017). Lignocellulosic biomass is readily available, cheap, and environment friendly (Yousuf et al., 2020). It is a versatile and renewable resource with numerous potential applications, including power generation, animal feed, biofuel generation, and the production of valuable products such as surfactants and bioplastics. Nonetheless, the efficient utilization of lignocellulosic biomass is challenging due to the differing chemical nature of its three major components, the strong and intricate linkages among them, and the overall recalcitrance of the biomass (Sahay, 2022). Typically, a pretreatment process is required to disrupt this recalcitrance, degrade lignin and hemicellulose, and reduce cellulose crystallinity. Available pretreatment technologies include physical, chemical, physiochemical, and biological methods.

One example of the lignocellulosic biomass from the agro-industrial side streams is palm kernel cake (PKC). After extracting oil from the kernels of oil palm fruits, the leftover pulp is formed into PKC. While palm kernel oil is used in commercial cooking and the formulation of soaps and detergents, the by-product PKC is used as high-protein feed for dairy cattle. Depending on the oil extraction techniques used, PKC can be classified as palm kernel meal (PKM) or palm kernel expeller (PKE) (Kini et al., 2020). PKM is generated from solvent extraction and contains less residual oil (< 3%), whereas PKE is produced from mechanical screw pressing and contains higher residual oil levels (8-10%). This gives PKE a greater energy density, making it a preferable choice over PKM in livestock production.

PKC is an attractive alternative to conventional feed ingredients like corn and soybean meal due to its low cost, year-round availability, and lack of competition with human food. However, its utilization efficiency is low in feeding non-ruminant livestock, mainly due to its high content of crude fiber or non-starch polysaccharides (NSP). NSPs consist of 78% galactomannan, 12% cellulose, 3% arabinoxylan and 3% glucuronoxylan (Düsterhöft et al., 1992). Not only mannan and cellulose are not readily digestible by monogastric animals, they are also responsible for protein encapsulation within the thick cell wall matrix (Gomez-Osorio et al., 2022). Consequently, the impaired nutrient digestibility has significantly limited the inclusion rate of PKC to no more than 20-25% in feeding non-ruminant livestock such as poultry, swine, and fish (Sharmila et al., 2014). In contrast, the recommended levels of PKC in the diets of goats and dairy cattle are as high as 30-50% (Sharmila et al., 2014). In commercial settings, PKE is typically applied at a much lower level, ranging from 3-5%, in feeding non-ruminant livestock. To increase the utilization efficiency of PKC in animal husbandry, it is necessary to improve nutrient digestibility, particularly by reducing the fibre content.

Fibre hydrolysis of PKC can be achieved through biological methods involving specific microbes or enzymes. Given the complex fibre matrix of lignocellulosic biomass, enzymatic pretreatment typically requires a large number and amount of enzymes, including accessory ones, making the process costly (Sahay, 2022). Moreover, the effect of enzymatic pretreatment is not always consistent across studies, likely due to variations in enzymes and diet formulations. In contrast, microbial fermentation offers a more economically favourable approach for transforming the bulk of low-value lignocellulosic biomass. Particularly, solid-state fermentation (SSF) by cellulolytic microorganisms has been extensively studied to improve the nutritional value of various agricultural by-products as functional components in animal husbandry. SSF involves the growth of microorganisms on moistened solid substrate in the absence of free-running water, enabling the use of a wide variety of agricultural waste as growth substrate and producing less liquid waste (Doriya et al., 2016; Webb, 2017). Fermenting microorganisms can secrete various extracellular enzymes to degrade polymeric substrates, improving nutrient digestibility. Additionally, the fermentation process can enhance the palatability and amino acid profile of the raw ingredient.

Numerous studies suggest that both bacterial and fungal fermentation could increase the total protein content and decrease the fibre content of PKC (Azizi et al., 2021). However, most studies have focused on the biotransformation of PKC involving extended fermentation times, which can increase production costs and elevate the risk of contamination. For instance, reduced crude fibre and increased crude protein contents were observed in PKC fermented with *Bacillus amyloliquefacien* and *Trichoderma harzianum* with a 7-day incubation period (Pasaribu et al., 2019). Similarly, Marzuki et al. demonstrated a significant increase in crude protein and decrease in crude fibre contents after fermenting PKE with *Aspergillus niger* for 66 h (Marzuki et al., 2008). Nonetheless, feeding rats with this fermented PKE as the sole protein source did not promote growth, possibly due to lower nitrogen digestibility of fungal protein and the presence of toxic secondary metabolites and undesirable odour.

In this study, we report the isolation of a high-mannanase producing strain of *Bacillus subtilis* capable of significantly decreasing the fibre content of PKC within a short fermentation period. Unlike most studies that focus on the fermentation of PKE, we have chosen to work on the fermentation of PKM, which has lower energy content. This makes PKM potentially more beneficial from the biotransformation process, leading to improved feed utilization. The first part of the study reports the isolation of *B. subtilis* for PKM fibre hydrolysis based on enzyme production capability when growing on the biomass. The effect on fibre degradation was assessed using neutral detergent fibre (NDF) analysis, measuring the total fibre in plant cell walls consisting of lignin, cellulose, and hemicellulose. Upon confirming its PKM-degrading capability, transcriptome analysis and liquid chromatography–mass spectrometry (LC-MS) were conducted to identify the enzymes involved in PKM fibre hydrolysis during the fermentation process. Two enzyme candidates were identified: β-mannanase GmuG and endoglucanase EglS. In the second part of the study, the enzyme candidates were expressed in *Escherichia coli* to obtain the purified recombinant enzymes for testing their PKM-degrading activity. The recombinant GmuG (rGmuG) demonstrated high fibre hydrolysis activity, releasing a substantial amount of reducing sugar from PKM as measured by dinitrosalicylic acid (DNS) assay. Additionally, the hydrolysis pattern of PKM fibre was elucidated through high-performance liquid chromatography (HPLC). Lastly, the biochemical properties of rGmuG were characterized in terms of optimum reaction temperature and pH, and substrate specificity. Our study provides an effective solution for improving the quality of PKM or other mannan-rich bioresources within a short fermentation time using the newly isolated *B. subtilis* strain.

## 2. Materials and methods

### 2.1 Sample and microorganisms

PKM was obtained from Wilmar palm kernel solvent extraction plant located in East Java, Indonesia. The bacteria populations used to inoculate PKM were obtained from exocarp of palm fruits from Wilmar’s oil palm plantations. The *B. subtilis* CK7 strain was obtained from PT. Wilmar Bernih Indonesia.

### 2.2 Isolation and screening of mannolytic bacteria

Microorganisms were collected from the exocarp of palm fruits in the form of palm fruit wash by rinsing the fruit surface with sterile water. About 0.1 – 0.5 ml of the palm fruit wash was inoculated into 1 g PKM in 5 ml sterile water for enrichment of PKM-degrading microbes. The PKM had been autoclaved at 121 °C for 15 mins prior to the enrichment process. After more than two weeks of incubation at 37°C, the grown bacteria were streaked on a Luria-Bertani (LB) agar plate to select for single colonies. Each bacteria colony was grown and re-inoculated into 100 ml M9 minimal medium (6.78 g/L Na_2_HPO_4_.7H_2_O, 3 g/L KH_2_PO_4_, 1 g/L NH_4_Cl, 0.5 g/L NaCl) containing 0.5% (w/v) locust bean gum (LBG) galactomannan (Sigma) as sole carbon source. The tubes were incubated at 37°C for 48 h and cell-free supernatant was collected as crude enzymes for the assessment of mannanase activity. Genomic DNA of isolates with high mannanase activity was extracted (Ausubel et al., 2003) and used as template in PCR for 16S rRNA partial gene amplification (~ 500 bp) using the following universal primers: 16S Forward (5’-CCTACGGGAGGCAGCAG-3’) and 16S Reverse (5’-GGACTACHVGGGTWTCTAAT-3’) (Takahashi et al., 2014). The amplicons were purified and sequenced. Sequence similarity and homology analysis against the GenBank database were carried out using the basic local alignment search tool (BLASTN) on the National Center for Biotechnology Information (NCBI) (https://www.ncbi.nlm.nih.gov/).

### 2.3 Palm kernel meal solid-state fermentation (SSF)

PKM was sterilized at 121°C for 20 min. In a 50 ml-tube, 1 g of PKM was inoculated with 6×10^8^ CFU of bacterial cells. The seed culture was prepared from the overnight liquid culture by centrifuging to harvest the cell pellet. This pellet was then resuspended in 1.5 ml of deionized water before inoculation into the PKM. SSF was carried out by incubating the inoculated PKM at 37°C for up to 24 h. After fermentation, 4 - 5 ml of deionized water was added to the fermented PKM, mixed with brief vortex, and centrifuged to get the cell-free supernatant for mannanase activity test. Alternatively, fermented PKM was dried at 60°C for fibre analysis.

### 2.4 Determination of mannanase activity

Mannanase activity was assayed by using a 0.5% (w/v) solution of LBG galactomannan in 100 mM sodium acetate buffer pH 5.0 (950 µl) to which 50 µl of the appropriately diluted enzyme solution was added. The reducing sugars released in 20 min or 60 min at 55°C at 1000 rpm were measured as mannose equivalents by the dinitrosalicylic acid (DNS) method described by Miller (Miller, 1959) with minor modifications. Briefly, 300 µl of the reaction mixture was reacted with 900 µl of DNS reagent at 100°C for 5 min followed by absorbance measurement at 540 nm. One unit of mannanase activity is defined as the amount of enzyme producing 1 µmol of mannose equivalents per min under the given conditions.

### 2.5 Determination of neutral detergent fibre (NDF) content

The control and fermented PKM was dried at 60°C and 0.5 g of the dried sample was used for NDF analysis according to the protocol based on AOAC 2003:04/ ISO 16472:2006. In brief, the sample was treated with sodium sulphite and neutral detergent solution using the default program in fibre analyzer (Fibertec 8000, FOSS). 1 L of neutral detergent solution with a final pH between 6.95 to 7.05 was prepared with 18.61 g of EDTA (disodium ethylene diamine tetraacetate), 6.81 g of sodium borate decahydrate, 30 g sodium lauryl sulphate, 10 ml of triethylene glycol and 4.56 g disodium hydrogen phosphate dissolved in water.

### 2.6 RNA extraction and transcriptome analysis

The overnight LB culture of *B. subtilis* F6 was centrifuged to yield a cell pellet of 12×10^8^ CFU, which was then directly used for RNA extraction to determine the RNA expression level before SSF. Another set of the cell pellet was resuspended in 3 ml sterile water and added to 2 g autoclaved PKM in a 50 ml tube for SSF at 37°C for 6 h. After fermentation, the fermented PKM was mixed with 8 ml phosphate-buffered saline (PBS) followed by filtration through miracloth (Sigma) to remove PKM debris and collect bacterial cells for RNA extraction. Three replicates were prepared for the samples before and after fermentation.

Total RNA was isolated using RNeasy Protect Bacteria Mini Kit (Qiagen, USA) including on-column DNase digestion. Quantifications and integrity were checked using RNA ScreenTape Assay with Agilent 4200 TapeStation System (Agilent). All samples had an RNA integrity number (RIN) > 9.

Subsequently, whole transcriptome sequencing was performed in order to examine the different gene expression profiles (Macrogen, South Korea). The library kit and type of sequencer were TruSeq Stranded Total RNA (NEB Microbe) and NovaSeq, respectively. Read mapping was performed with *B. subtilis* BSn5 as the reference strain. Transcript expression level was quantified in term of RPKM (Reads Per Kilobase of transcript per Million mapped reads) values. The differential expression of a specific gene was quantified in terms of fold change of RPKM values between post- and pre-fermentation, presented in logarithmic scale (base 2). A positive log2 fold change indicates increased expression, while a negative value indicates decreased expression.

### 2.7 Genomic DNA preparation and whole genome sequencing

The genomic DNA was isolated according to Hoffman (Ausubel et al., 2003) with minor modifications. Briefly, the pellet of bacterial cells collected from the overnight culture was resuspended in 200 µl breaking buffer (2% (v/v) Triton X-100; 1% (v/v) sodium dodecyl sulphate (SDS); 100 mM NaCl; 10 mM Tris-HCl, pH 8.0; 1 mM EDTA, pH 8.0). Subsequently, about 0.3 g glass beads (~200 µl volume), 200 µl phenol/chloroform/isoamyl alcohol and 200 µl TE buffer were added. Next, microcentrifugation was performed for 5 min at 12,000 rpm and the aqueous layer was transferred to a clean microcentrifuge tube. 1 ml of absolute ethanol was added and mixed by inversion. Microcentrifugation was performed for 3 min at 12,000 rpm and the pellet was resuspended in 0.4 ml TE buffer. A total of 30 µl of 1mg/ml DNase-free RNase A was added, mixed, and incubated for 5 min at 37°C. Then, 10 µl of 4 M ammonium acetate and 1 ml of absolute ethanol were added and mixed by inversion. Microcentrifugation was performed for 3 min at 12,000 rpm and the dry pellet was resuspended in 100 µl TE buffer. The integrity and quality of the DNA was assessed by 0.8 % agarose gel electrophoresis and Nanodrop spectrophotometer determining A260/280 ratio, respectively. The DNA was sent to NovogeneAIT for microbial whole genome sequencing with Illumina HiSeq PE150 sequencing platform.

### 2.8 Expression of recombinant proteins GmuG and EglS

Gene fragments of *gmuG* and *eglS* were amplified from the purified genome of *B. subtilis* F6 and inserted into pET-28a(+) plasmid. Following plasmid construction in the cloning strain of *E. coli* DH5α, the plasmids were extracted using AxyPrep™ Plasmid Miniprep Kit (Axygen, Corning, USA). After the plasmid sequence was verified, the purified plasmids were introduced into the protein expression strain of *E. coli* Rosetta. The culture was induced for 4 h at 37 °C by adding 0.5 mM isopropyl-β-d-thiogalactopyranoside (IPTG). The culture was then harvested by centrifugation at 6500 rpm, 15 min, 4°C and the cell pellet was frozen at −20 °C.

### 2.9 Purification of GmuG and EglS recombinant proteins

The cell pellet was resuspended in BugBuster (primary amine-free) extraction reagent (Sigma) supplemented with lysonase. The supernatant was then loaded onto a Ni NTA Beads 6FF (Bio Basic) equilibrated in 50 mM NaH_2_PO_4_, 300 mM NaCl, and 10 mM imidazole, pH 8.0, at room temperature. The resin was then washed with 50 mM NaH_2_PO_4_, 300 mM NaCl, and 20 mM imidazole, pH 8.0. Elution was accomplished with 50 mM NaH_2_PO_4_, 300 mM NaCl, and 500 mM imidazole, pH 8.0.

### 2.10 SDS-PAGE analysis

Samples were resolved either on Mini-PROTEAN TGX Stain-Free Precast Gels or Mini-PROTEAN TGX Gels (4-20%) (Biorad, USA) in running buffer. For the latter, the proteins were stained with Coomassie brilliant blue.

### 2.11 PKM hydrolysis experiments

The hydrolysis of PKM was carried out at 37°C statically in an incubator. Each reaction consisted of 2 g autoclaved PKM submerged in 10 ml enzyme solution. Enzyme solution was prepared in distilled water and the protein concentration was determined using Bradford reagent (VWR, USA) with bovine serum albumin as the protein standard. In the first experiment, purified enzyme (rGmuG, rEglS) or commercial β-mannanase (Winovazyme Biotech, China) was added at an equal molarity of 3 nmol, which is approximately equivalent to 120 µg of GmuG or commercial β-mannanase, or 165 µg of EglS. In the second experiment, the enzyme was added at an equal mass of 120 µg while the mixed-enzyme treatment group consisted of 60 µg of rGmuG and rEglS each. Aliquots of PKM hydrolysate (150 µl) were removed at different timepoints, centrifuged (12,000 rpm, 1 min), and the amount of reducing sugars in the supernatant was determined using DNS method as described previously.

### 2.12 High performance liquid chromatography (HPLC) for sugar analysis

The sugar determination was done using HPLC with the refractive index detector (RID). The chromatography was performed with Agilent 1260 Infinity II, installed with Agilent Hi-Plex Na guard column (7.7 x 50 mm). Separation of sugar monomers and oligosaccharides was achieved using Agilent Hi-Plex Na analytical column (7.7 x 300 mm) in 45 min with an isocratic flow of 0.3 ml/ min using 100% Milli-Q water. Samples were injected using an auto-sampler and the injection volume was 5 μl. Column temperature was 85°C. The standards mannohexaose, mannopentaose, mannotetraose, mannotriose and mannobiose were purchased from Megazyme (Ireland). Mannose, cellobiose, xylose and galactose were purchased from Sigma-Aldrich (Germany). Glucose was purchased from Bio Basic (Singapore).

### 2.13 Determination of optimum pH and optimum temperature

The effects of pH and temperature on the purified rGmuG activity were evaluated under standard reaction condition as described above using 0.5 % (w/v) LBG as substrate. The optimum pH of rMan3 was determined at 55°C in four different buffer systems (each 100 mM): glycine-HCl buffer (pH 3.0), sodium acetate buffer (pH 4.0 - 5.0), Tris-HCl buffer (pH 6.0 - 8.0), and glycine-NaOH buffer (pH 9.0 – 10.0). The optimum temperature was investigated in 100 mM sodium acetate buffer (pH 5.0) containing 0.5 % (w/v) LBG as substrate over a range of temperature (30°C to 70°C).

### 2.14 Determination of substrate specificity

The activity of the purified rGmuG was determined at 55°C, 1000 rpm in 20 min on 0.5 % (w/v) substrates: 1,4-β-D-mannan (Megazyme) (pH 5.0), LBG galactomannan (Sigma) (pH 5.0), konjac glucomannan (Megazyme) (pH 5.0), xylan (from beechwood) (Carlroth) (pH 4.97), sodium carboxymethyl cellulose (Sigma) (pH 5.01) and polygalacturonic acid (Sigma) (pH 4.49).

## 3 Results and discussion

### 3.1 Isolation of B. subtilis F6 from palm fruit exocarp

In this study, microbial community from the exocarps of oil palm fruits was collected as the source of microbes and incubated with sterilized PKM for enrichment of PKM-degrading microbes. The high abundance of mannan fibre would likely help enrich microbes capable of utilizing mannan as energy source for growth. Bacterial isolates enriched by long-hour PKM incubation were isolated and examined for their mannanase activity before re-introduced into PKM for further incubation **(Figure 1)**. Among the mannanase-producing bacterial isolates, F6 was one of the strains that consistently showed high mannanase activity. The strain was identified as *B. subtilis* based on >98% pairwise identity upon comparison of the 16S RNA sequence with database from GenBank. Species such as *Bacillus*, *Enterococcus*, and *Lactobacillus* have been recognized for their probiotic applications in animal feed (Varzakas et al., 2018). Not only *B. subtilis* has potential probiotic properties, but it is also considered as GRAS (Generally Regarded as Safe) organism by the Food and Drug Administration (FDA).

**Figure 1.**
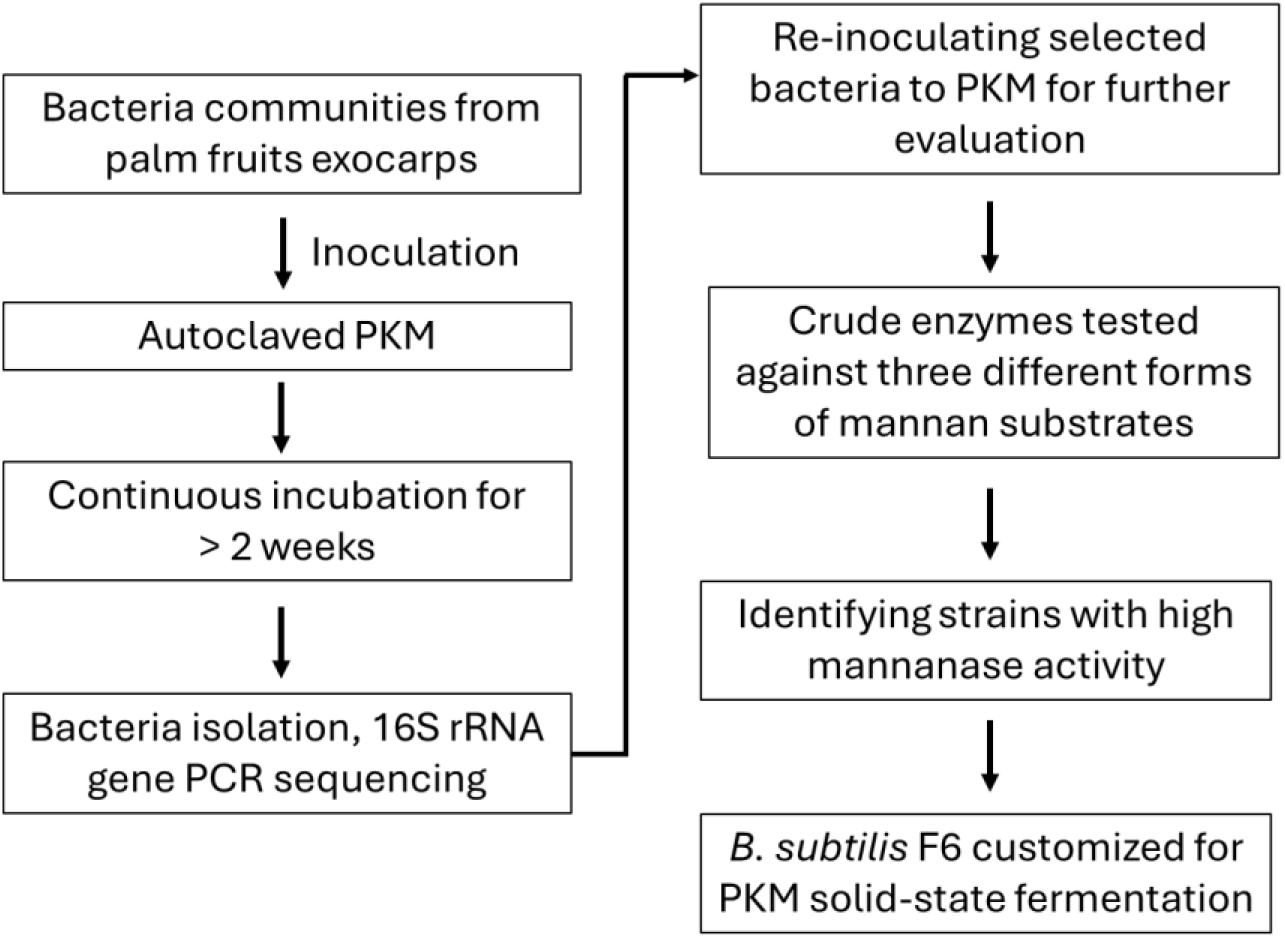
Screening process for isolating beneficial bacteria for PKM SSF. *B. subtilis* strain F6 was selected based on its secreted mannanase activity upon incubation with near solid-state PKM.

While fermenting microorganisms can be obtained from cell collection centres, our focus on undomesticated environmental isolates taps into the abundant and diverse microbial biodiversity in nature. *B. subtilis* F6 was able to thrive on near solid-state PKM in our initial screening process over long incubation period, secreting high mannanase activity. During the extended incubation period, it is presumed that readily available carbon sources, such as residual oil or simple sugars, were first utilized by bacterial cells before they began producing hydrolytic enzymes to degrade the complex polysaccharides. Furthermore, the prolonged incubation over several weeks facilitated the enrichment of microbes capable of thriving on PKM, eventually leading to their dominance within the microbial community.

Previously, Sari et. al. isolated cellulolytic and mannolytic aerobic bacteria from buffalo rumen for PKM fibre degradation (Sari et al., 2021). The bacteria were isolated following enrichment in a liquid medium composed of mineral salt solution, yeast extract and 1% PKM. In our study, the PKM-degrading microbes were isolated from an extended incubation of the bacteria with PKM fibre. We applied stringent growth conditions, enriching microbes in an environment closer to PKM SSF conditions without supplementation of additional nutrients, thus facilitating the isolation of strains that can thrive on near solid-state PKM. On the other hand, Virginia et al. isolated a mannanase-producing *B. subtilis* strain CK7 from a palm oil mill area and the mannanase was demonstrated its potential for hydrolysing PKE (Virginia et al., 2018). However, the direct fibre-degrading capability of the CK7 strain under PKM SSF conditions remains unexplored.

### 3.2 Comparative analysis of mannanase activity between F6 and CK7 strains

Strain-level variation is a basic feature of bacteria that dictates their survival in diverse environmental niches and is a key factor in determining their physiology and other traits (Nie et al., 2022). It is known that strains with almost identical genomes can exhibit differing physiological characteristics. The variability in the genome between individual strains may be small and well defined, but it may cause large phenotypic changes (e.g. point mutations causing drug resistance) (Van Helden, 1998). In this study, the newly isolated strain of *B. subtilis* F6 was compared with another mannanase-producing environmental isolate CK7 for their mannanase production under PKM SSF conditions over 24 h, using an equal amount of initial inoculum (6×10^8^ CFU/g of PKM). Mannanase activity was assessed at different fermentation time points using three distinct mannan-based substrates: LBG galactomannan, konjac glucomannan, and 1,4-β-D-mannan **(Figure 2A-C)**. Substantially higher level of mannanase activity across different timepoints was observed with F6 strain compared to the CK7 strain. As early as 12 h, strong mannanase activity was detected in PKM fermented with F6 strain, and the enzymatic activity was consistently enriched throughout the sampling times. This indicates a relatively fast mannanase production in response to growing on PKM. Among the different mannan-based substrates, the crude enzyme produced by F6 strain showed the highest activity with LBG **(Figure 2A)**, followed by konjac **(Figure 2B)** and 1,4-β-D-mannan **(Figure 2C)**, indicating the differential substrate preference of the enzyme. In contrast, the CK7 strain did not respond well upon contact with solid-state PKM despite its previously reported LBG-inducible mannanase activity in liquid medium (Virginia et al., 2018). The variance in mannanase activity between the two strains may stem from differences in their adaptability and physiological characteristics when cultivated on solid-state PKM. The newly isolated F6 strain demonstrated notable responsiveness to PKM, swiftly initiating the production of substantial mannanase activity within a short period of time.

**Figure 2.**
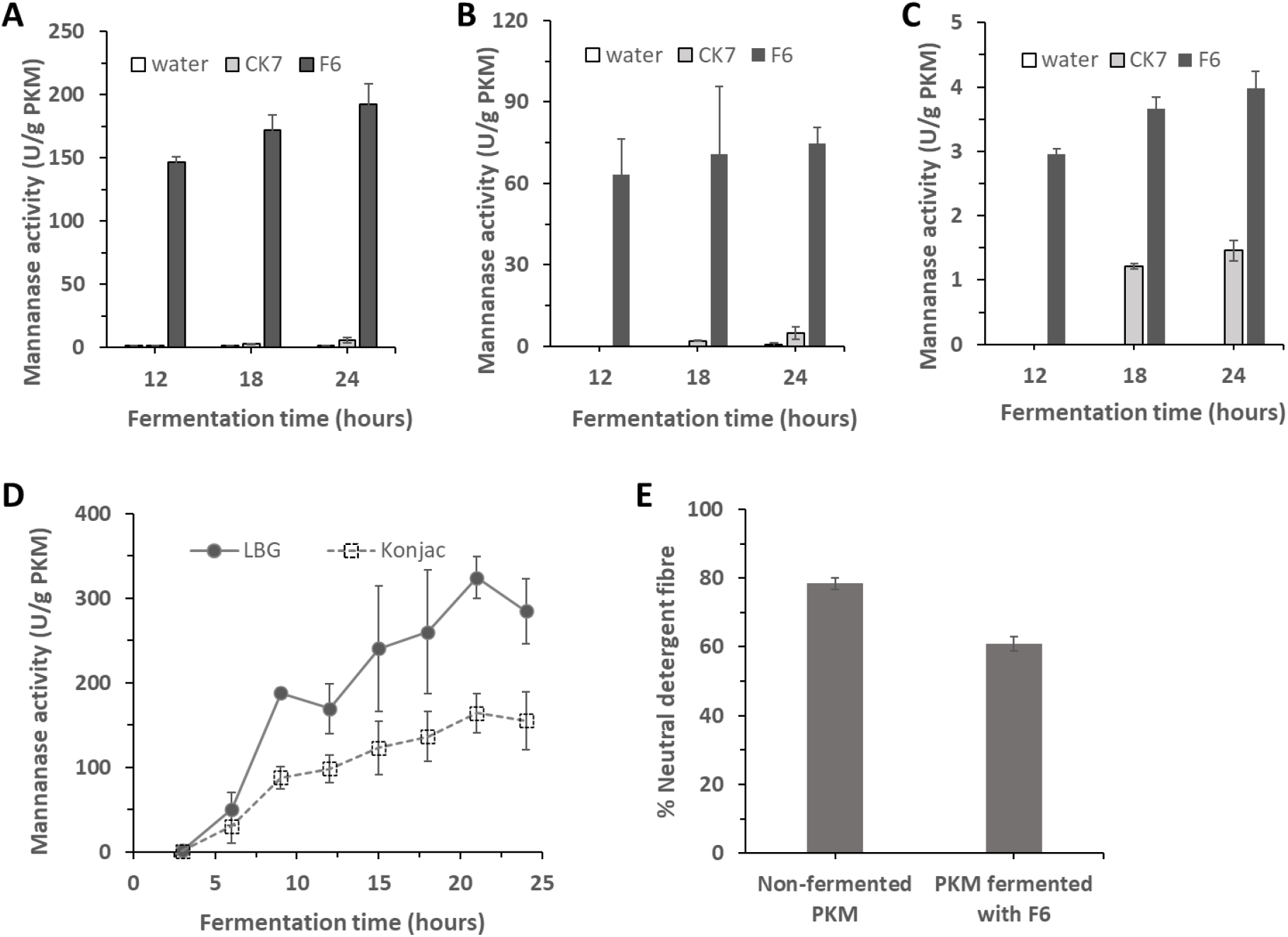
Evaluating the mannanase activity and fibre hydrolysis of *B. subtilis* strain F6. (A – C) Performance comparison between *B. subtilis* F6 and CK7 in PKM SSF. The isolated strain F6 showed significant enhancement of enzymatic hydrolysis activity against 3 different forms of mannan-based substrates at three different fermentation times: (A) LBG galactomannan, (B) konjac glucomannan, and (C) 1,4-β-D-mannan, as compared to strain CK7. (D) Time-dependent performance of strain F6 during PKM SSF assessed by its enzymatic hydrolysis activity against LBG and konjac. The mannanase activity assay was performed at 55°C for 60 min. The reaction comprised of 0.5% (w/v) substrate in 0.1 M sodium acetate buffer, pH 5.0. (E) Neutral detergent fibre (NDF) content of non-fermented PKM and PKM fermented with *B. subtilis* F6. Data are the mean of three replicates, and bars indicate standard deviation of three replicates.

### 3.3 Mannanase production and fibre hydrolysis during PKM SSF

The F6 strain was further examined for their mannanase production at 3 h-intervals across the PKM SSF process. As early as 6 h, considerable mannanase activity was detected and a sustained increase was observed in the following timepoints **(Figure 2D)**. The enzyme activity reached the plateau phase within 24 h of fermentation. The effect of mannanase production was subsequently examined on the extent of PKM fibre hydrolysis by measuring the residual neutral detergent fibre (NDF) content after fermentation. The NDF method is used for the determination of total insoluble fibres in food and feed by estimating of the content of hemicellulose, cellulose and lignin in the sample. As shown in **Figure 2E**, the NDF level significantly decreased from 78.4% to 60.9% after an overnight fermentation with F6 strain.

The digestibility of PKM for feeding non-ruminant livestock can be enhanced through reducing the fibre content. Previously, Marzuki et al. reported a substantial decrease of NDF content from 79.0% to 50.3% in PKE fermented with *A. niger* for 66 h (Marzuki et al., 2008). Although a higher level of fibre degradation was attained in the *A. niger*-fermented PKE, the extended fermentation duration necessary for this process may not be economically favourable due to increased production costs and potential reductions in overall productivity. Hence, the newly isolated *B.* strain provides an alternative for achieving substantial fibre degradation within a shorter time. Given the advancements in synthetic biology, the F6 strain can be further engineered to enhance its fibre hydrolysis capacity without necessitating prolonged fermentation periods for fibre degradation.

Overall, the results suggest that enzymes produced by the newly isolated F6 strain are effective in breaking down the PKM fibre. The enzymatic breakdown of PKM fibre likely involves a combination of various hydrolases, such as mannanase, cellulase, and xylanase. This complexity arises from PKM’s composition, which comprises different NSPs, with mannan being the predominant fibre component. Understanding the enzymes involved in PKM fibre hydrolysis would provide mechanistic insights into fibre reduction during PKM fermentation with the F6 strain. To identify the enzyme(s) responsible for PKM fibre degradation, it is necessary to study the expression levels of different hydrolases on a whole-genome level, which can be efficiently accomplished through transcriptome analysis.

### 3.4 Analysis of hydrolase expression level and secretion

Bacterial strains growing in different environmental conditions can be molecularly characterized. In this study, transcriptome analysis was used to identify the genes involved in PKM fibre degradation along with other metabolic responses in the F6 strain cultivated on solid-state PKM. RNA was isolated from the bacterial cells at the end of 6-h PKM SSF, when a significant level of mannanase activity can be detected. Gene expression levels were compared to that of the bacterial cells grown in liquid LB culture overnight.

Among the genes coding for enzymes responsible for the degradation of NSPs, upregulations in the expression level were observed in mannanase, endoglucanase, xylanase and pectate lyase **(Table 1A)**. The highest fold change was observed with mannanase *gmuG* (45.17-fold), followed by pectate lyase (7.12-fold) and endoglucanase *eglS* (6.09-fold). Interestingly, there are two distinct mannanase genes in the F6 strain, sharing a pairwise identity of 67.9%. However, only the *gmuG* gene showed an upregulation while another gene showed a downregulation (0.16-fold) in the expression level. This suggests that the two genes are likely associated with different metabolic or physiological functions. The *gmuG* gene of *B. subtilis* is located within the glucomannanan utilization operon (*gmuBACDREFG*), in which all the genes were upregulated in their expression level after 6h-PKM SSF **(Table 1B)**. β-mannanase encoded by *gmuG* is secreted extracellularly to hydrolyse mannan into mannooligosaccharides which are then transported into the *Bacillus* cells by phosphotransferase system made up of GmuA, B and C (Sadaie et al., 2008). Within the cells, the oligosaccharides are further processed by β-glucosidase (GmuD), fructokinase (GmuE), or isomerase (GmuF) for subsequent metabolic processes. While the operon is induced by the degradation products of glucomannan such as mannobiose, it can be repressed by an internal repressor (GmuR) located within the operon and glucose via the carbon catabolite repression system using CcpA.

**Table 1.** Variation of gene expression in *B. subtilis* F6 before and after PKM solid-state fermentation (SSF)

| Gene name | Expression before SSF (RPKM) | Expression after SSF (RPKM) | Log<sub>2</sub> (fold change) After SSF vs Before SSF |
| --- | --- | --- | --- |
| **A. Enzymes involved in NSP degradation** |  |  |  |
| Mannanase (*gmuG*) | 7.3 ± 2.2 | 329.9 ± 45.3 | 5.5 |
| Pectate lyase 2 | 38.5 ± 1.3 | 274.1 ± 44.3 | 2.8 |
| Endoglucanase (*eglS*) | 32.7 ± 8.0 | 198.9 ± 93.4 | 2.6 |
| Xylanase (*xynA*) | 29.0 ± 7.4 | 155.7 ± 33.1 | 2.4 |
| Xylanase (*xynC*) | 115.8 ± 30.0 | 419.7 ± 142.7 | 1.9 |
| Pectate lyase 1 | 905.1 ± 291.6 | 1883.5 ± 472.8 | 1.1 |
| β-glucanase (*bglS*) | 145.1 ± 72.4 | 100.4 ± 9.0 | -0.5 |
| Mannanase | 12.3 ± 1.8 | 2.0 ± 0.5 | -2.6 |
| **B. Glucomannan utilization operon** |  |  |  |
| PTS sugar transporter subunit IIB (*gmuB*) | 4.9 ± 3.3 | 331.2 ± 86.2 | 6.1 |
| Phosphotransferase enzyme IIA (*gmuA*) | 5.2 ± 2.7 | 197.7 ± 56.4 | 5.2 |
| PTS system cellobiose-specific IIC (*gmuC*) | 2.6 ± 0.5 | 339.6 ± 87.2 | 7.0 |
| Glycoside hydrolase family 1 protein (*gmuD*) | 5.8 ± 0.7 | 623.5 ± 143.8 | 6.7 |
| GntR family transcriptional regulator (*gmuR*) | 20.5 ± 9.6 | 1433.1 ± 359.4 | 6.1 |
| Fructokinase (*gmuE*) | 1.7 ± 0.6 | 210.6 ± 38.3 | 7.0 |
| Mannose-6-phosphate isomerase (*gmuF*) | 10.2 ± 3.4 | 874.3 ± 148.5 | 6.4 |
| β-mannosidase (*gmuG*) | 7.3 ± 2.2 | 329.9 ± 45.3 | 5.5 |
| **C. Mannose operon** |  |  |  |
| PTS mannose transporter subunit IIABC (*manP*) | 0.8 ± 0.5 | 12.7 ± 10.8 | 4.0 |
| Mannose-6-phosphate isomerase, class I (*manA*) | 3.1 ± 0.6 | 45.0 ± 34.1 | 3.9 |
| YjdF family protein (*yjdF*) | 2.7 ± 2.3 | 27.6 ± 13.6 | 3.4 |
| **D. Sec translocase components** |  |  |  |
| Protein translocase subunit (*secDF*) | 83.6 ± 6.9 | 314.3 ± 24.4 | 1.9 |
| Preprotein translocase subunit (*secG*) | 649.8 ± 314.2 | 1613.6 ± 65.2 | 1.3 |
| Preprotein translocase subunit (*secA*) | 150.0 ± 13.5 | 638.6 ± 38.9 | 2.1 |
| Preprotein translocase subunit (*secE*) | 7.4 ± 7.9 | 331.2 ± 33.9 | 5.5 |
| Preprotein translocase subunit (*secY*) | 1361.6 ± 256.5 | 7646.3 ± 1092.2 | 2.5 |

Meanwhile, another cluster of genes denoted as mannose operon, comprising *manP*, *manA* and *yjdF*, had their expression level upregulated as well **(Table 1C)**. Nonetheless, their expression level and the respective fold change were much lower than that of glucomannan utilization operon. For instance, mannose-6-phosphate isomerases (MPI), which convert mannose-6-phosphate into fructose-6-phosphate entering the glycolysis pathway, are encoded by *gmuF* and *manA* in glucomannan utilization operon and mannose operon, respectively. While *gmuF* was expressed at 874 RPKM with a fold change of 85.9, *manA* was expressed at 45 RPKM with a fold change of 14.5 after 6 h-PKM SSF. The results suggest that glucomannan utilization operon is likely the dominant gene cluster involved in mannose metabolism under the PKM SSF conditions.

In addition, a significant upregulation of the expression level was observed in Sec translocase components **(Table 1D)**. Being the main pathway for protein secretion in *B. subtilis*, an upregulated Sec-mediated protein secretion would promote the secretion of various hydrolytic enzymes necessary for the breakdown of macromolecules, facilitating nutrient uptake during fermentation. Furthermore, the fermentation conditions also seemed to favour flagellar assembly, sporulation, biofilm formation and antibiotics biosynthesis.

The observed changes in the expression levels were further verified by the detection of secretory proteins in fermented PKM. To shortlist the candidate enzymes likely involved in PKM fibre degradation, water-soluble proteins were extracted from the fermented PKM and analysed using LC-MS after trypsin digestion. Carbohydrases detected with the highest number of unique peptide and highest percentage of sequence coverage are β-mannanase GmuG and endoglucanase EglS, which also had their gene expression levels upregulated according to the transcriptome analysis performed earlier. Considering that both mannan and cellulose are part of the indigestible components of PKM fibre, with mannan being the primary component, the two enzymes GmuG and EglS warrant a further investigation for their respective role in PKM fibre degradation. Apart from carbohydrases, other proteins detected in the fermented PKM include extracellular neutral metalloprotease NprE and peptidase S8 (subtilisin family). During post-exponential growth, *B. subtilis* secretes among other enzymes several proteases (L.-F. Wang et al., 1989). AprE (subtilisin) and NprE are the most abundant proteases and are found in the culture medium during stationary phase where they contribute >95% of the extracellular proteolytic activity of *B. subtilis* (Harwood & Kikuchi, 2022).

### 3.5 Protein overexpression in E. coli to obtain purified enzymes

Alignment of the amino acid sequence of GmuG and EglS between *B. subtilis* F6 and reference strain 168 showed an identity of 99.4% and 99.8%, respectively. Given the high identity of protein sequence, the protein features of GmuG and EglS derived from strain F6 were predicted with reference to that of strain 168 **(Figure 3A)**. To further investigate the role of GmuG and EglS in PKM fibre degradation, purified recombinant enzymes were first obtained by protein overexpression in *E. coli*. Using the genomic DNA of *B. subtilis* F6 as template, *gmuG* and *eglS* genes were amplified by PCR and cloned into a pET-28a (+) plasmid for expression in *E. coli*. **Figure 3B** shows the successful expression of the recombinant proteins in *E. coli* Rosetta. Following His-tag purification, purified recombinant proteins of GmuG (rGmuG) and EglS (rEglS) were obtained **(Figure 3C)**.

**Figure 3.**
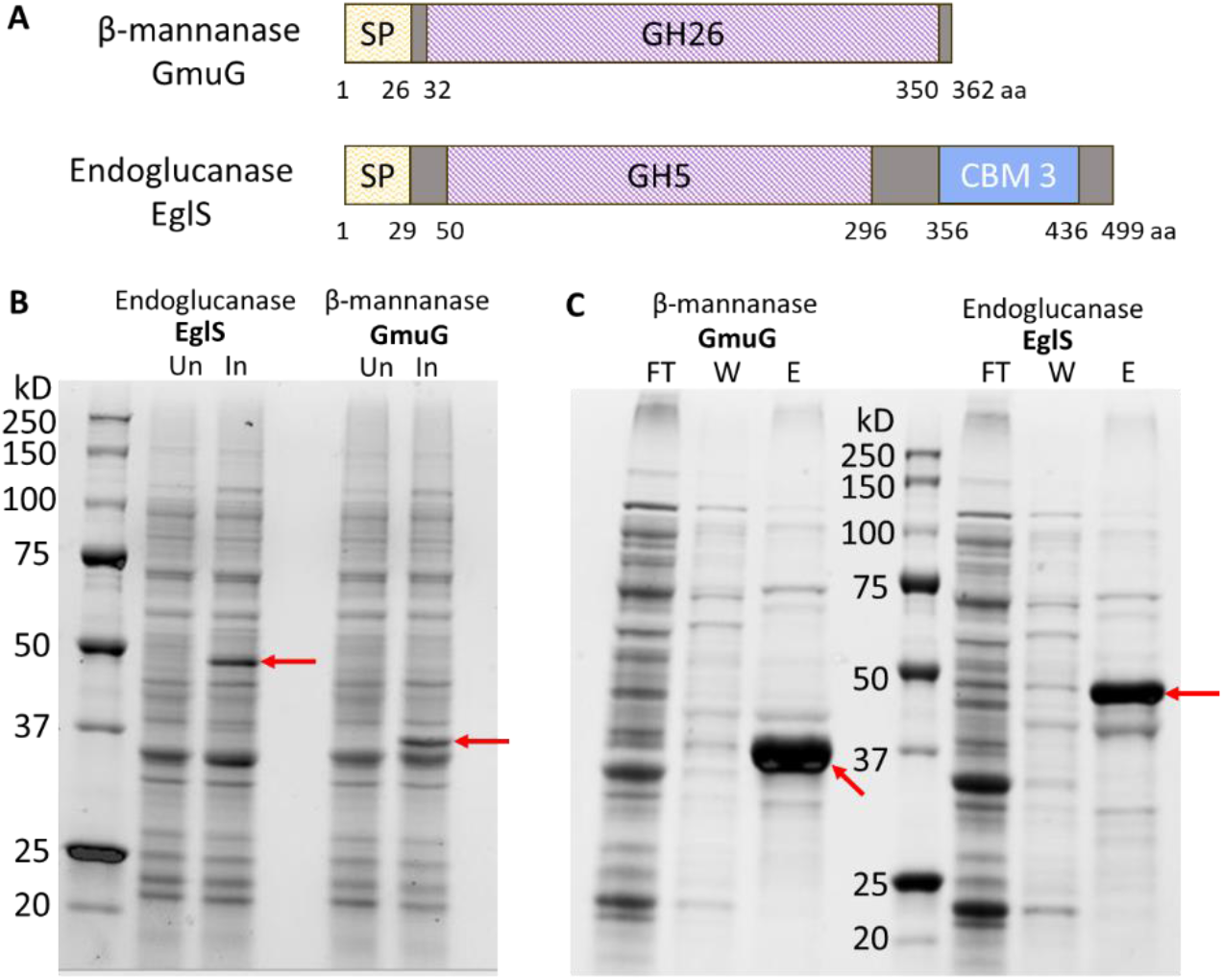
Overexpression of F6-derived β-mannanase GmuG and endoglucanase EglS in *E. coli* to get purified proteins. (A) Protein domains of F6-derived GmuG and EglS predicted based on the proteins from *B. subtilis* reference strain 168. GmuG preprotein has 362 amino acids, with an N-terminal signal peptide and a GH26 catalytic domain. The deduced mature protein contains 336 residues with a calculated mass of 37.0 kDa. EglS preprotein has 499 amino acids, containing an N-terminal signal peptide, a GH5 catalytic domain and a CBM 3. The deduced mature protein has 470 amino acids with a calculated protein size of 51.7 kDa. SP: signal peptide; GH: glycoside hydrolase; CBM: carbohydrate-binding module; aa: amino acid. (B, C) SDS-PAGE analysis of protein expression in whole cell lysate (B) after induction and (C) after His-tag purification. The bands corresponding to GmuG or EglS were marked. Un: Uninduced; In: Induced; FT: flow-through; W: wash; E: elution.

### 3.6 PKM hydrolysis using purified enzymes rGmuG and rEglS

Purified enzymes rGmuG and rEglS were added to the PKM at an equal molarity of 3 nmol **(Figure 4A)** or an equal mass of 120 µg **(Figure 4B)** in two different reactions. PKM samples were examined for the amount of reducing sugar released across different timepoints using colorimetric DNS assay. A gradual release of reducing sugar from PKM treated with mannanase rGmuG was observed over time. A comparable hydrolysis activity was also observed between rGmuG and the commercial β-mannanase under the conditions tested. In contrast, endoglucanase rEglS was not able to release any reducing sugars from PKM. To test the synergistic effect of rGmuG and rEglS on PKM fibre hydrolysis, PKM was treated with an enzyme mix of the two hydrolases at 60 µg each **(Figure 4B)**. However, the enzyme mix did not perform any better than using 120 µg of rGmuG alone. The results suggest that mannanase rGmuG was effective in breaking down the PKM fibre and likely responsible for the PKM fibre degradation as observed earlier on during the SSF process. On the other hand, endoglucanase rEglS neither showed hydrolysis activity towards PKM fibre on its own nor exhibited synergistic effect with enzyme rGmuG in PKM fibre degradation. It could be due to the problem with substrate specificity or accessibility. Particularly, as mannan is the most abundant NSP in PKM, cellulose could be so tightly entangled with the bulk of mannan fibre that accessibility of cellulose for hydrolysis is highly limited when the mannan is not degraded enough.

**Figure 4.**
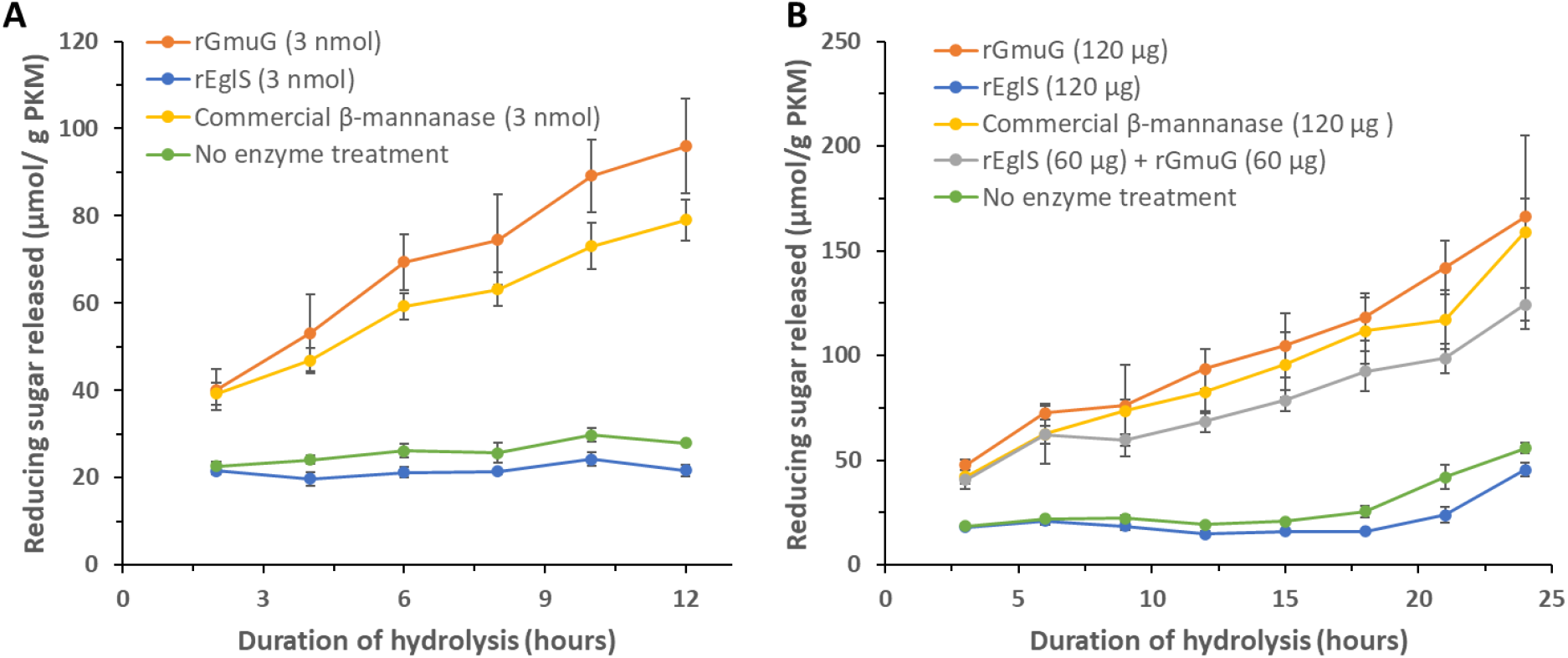
PKM hydrolysis experiments. Fibre hydrolysis activity of the enzymes was estimated by measuring the amount of reducing sugars released from PKM. The enzymes tested were purified recombinant enzymes, β-mannanase rGmuG and endoglucanase rEglS, sourced from *B. subtilis* F6, and commercial β-mannanase from Winovazyme. Reaction (A) consisted of 2 g PKM and 3 nmol enzyme in 10 ml distilled water. Reaction (B) consisted of 2 g PKM and 120 µg enzyme in 10 ml distilled water. Reaction was performed at 37°C statically. Aliquots were withdrawn at different timepoints for analysis. Data are the mean of three replicates, and bars indicate standard deviation of three replicates.

Next, the hydrolysis activity of mannanase rGmuG was further characterized in term of its hydrolysis pattern of PKM fibre using HPLC analysis. While the DNS method provides a non-specific way to estimate the concentration of reducing sugars in enzymatic hydrolysates, a more in-depth analysis can be done using HPLC for the identification and quantification of individual sugars in the complex sugar mixture derived from lignocellulosic biomass. Agilent Hi-Plex Na sugar column enabled the separation of mannan and cellulose degradation products of different sizes. Unfortunately, mannose, galactose, and xylose could not be distinguished due to their similar retention time of 34 minutes in the column. Given that mannan is the predominant NSP in PKM, the peak was inferred to primarily comprise its degradation product, mannose. Water-soluble sugars were extracted from PKM samples, with or without enzymatic treatment using mannanase rGmuG for 22 h. The control PKM sample without enzymatic treatment exhibited a moderate level of cellobiose and low amount of mannose and glucose **(Figure 5)**. PKM fibre hydrolysis by rGmuG resulted in the release of mainly mannobiose and mannotriose, along with small traces of mannose and other larger mannooligosaccharides. Notably, there was no increase in the levels of cellulose degradation products such as glucose and cellobiose, indicating no cross-reactivity of rGmuG towards cellulose in PKM. Overall, our results demonstrate the activity of mannanase rGmuG in breaking down PKM fibre, releasing mannose and mannooligosaccharides.

**Figure 5.**
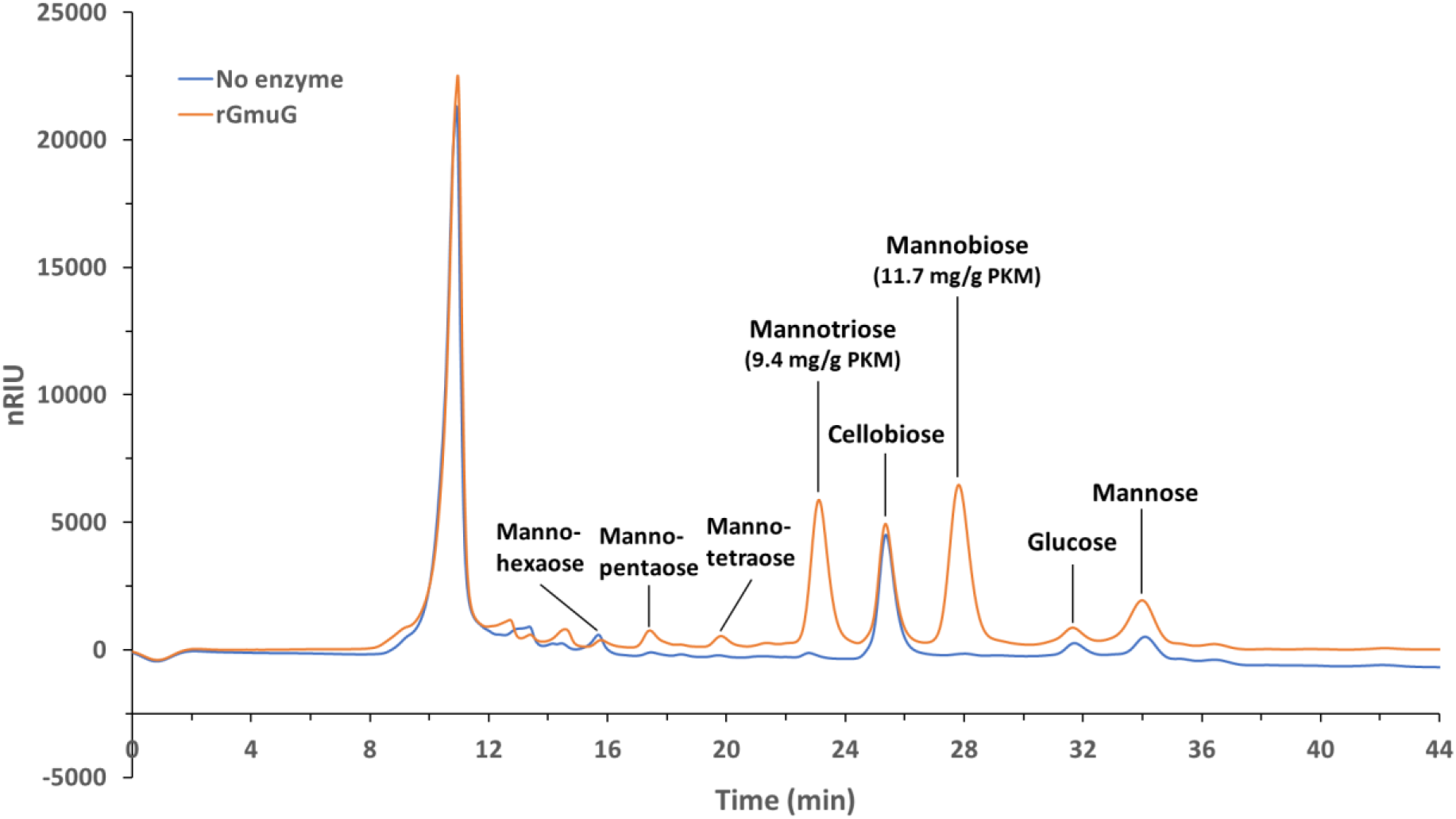
HPLC chromatogram showing the sugar profile of PKM samples treated with (orange) or without (blue) enzyme rGmuG. 1 g autoclaved PKM was treated with 50 µg enzyme and topped up with distilled water to a total water content of 5 ml. The enzymatic treatment was performed at 37°C, 220 rpm for 22 h. The untreated and treated PKM samples had a total releasing sugar of 12.2 and 36.3 mg/g PKM, respectively.

The hydrolysis pattern of rGmuG is similar to the mannanases reported previously. For instance, Jana & Kango reported the release of mannose, mannobiose and mannotriose from PKC treated with β-mannanase purified from *Aspergillus oryzae* (Jana & Kango, 2020). Besides, the extensive hydrolysis of ivory nut mannan by the purified mannanase from *Sclerotium rolfsii* yielded mainly mannobiose and mannotriose (Sachslehner & Haltrich, 1999). Given that small amounts of mannose were also generated by rGmuG, the enzyme seems to exhibit both endo- and exo-acting hydrolysis activities towards the mannan in PKM. Although most endo-1,4-β-mannanases hydrolyse mannan polysaccharides to produce mainly mannobiose and mannotriose with no free mannose (Liao et al., 2014), several studies also observed the release of mannose by endo-β-mannanases (Luo et al., 2009; Y. Wang et al., 2012). In fact, mannanases belong to the GH26 family were reported to have endo-β-1,4-mannanase, exo-β-1,4-mannobiohydrolase or mannobiose-producing exo-β-mannanase activities. Cervero et al. reported that free mannose was released upon hydrolysis of 5% PKC using 10% (v/w) of commercial endo-mannanase Mannaway derived from a *Bacillus* strain. They proposed that Mannaway had the capability to act on even small oligosaccharides or close to chain ends to release free mannose (Cerveró et al., 2010). On the other hand, recombinant endo-mannanase rPoMan5A from *Penicillium oxalicum* GZ-2 was able to release mannose along with other mannooligosaccharides from konjac glucomannan. They proposed that rPoMan5A functions as both endo-β-1,4-mannanase and 1,4-β-mannosidase (Liao et al., 2014).

### 3.7 Characterization of rGmuG biochemical properties

Following the identification of mannanase GmuG as the primary catalyst responsible for PKM fibre degradation, a more detailed understanding of the enzyme was attained by investigating its biochemical properties. This included determining its optimum reaction temperature and pH, and substrate specificity. Using LBG galactomannan as substrate, rGmuG showed the highest activity in the temperature range of 50 - 55°C **(Figure 6A)**. Interestingly, rGmuG exhibited two pH optima at around 5.0 and 9.0 **(Figure 6B)**. Below pH 5.0, the enzyme activity reduced drastically till there was no activity at pH 4.0.

**Figure 6.**
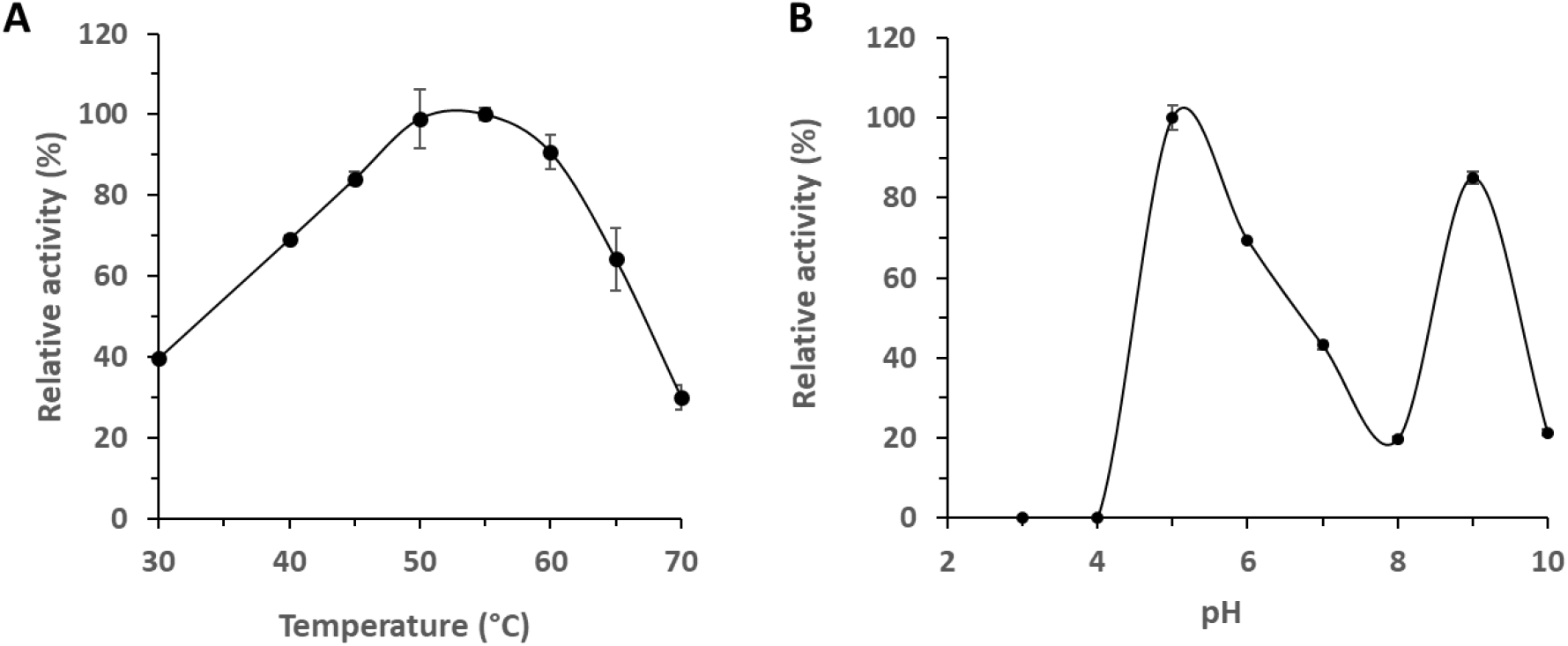
Biochemical properties of the purified β-mannanase rGmuG. (A) Optimum temperature and (B) pH of rGmuG. Effect of temperature on enzyme activity was measured at different temperatures using 0.5% LBG as substrate. The influence of pH on enzyme activity was determined in different 0.1 M buffers using 0.5% LBG as substrate. Data are the mean of three replicates, and bars indicate standard deviation of three replicates.

Most endo-β-mannanases showed maximal activity in the temperature range of 40 to 65°C (Srivastava & Kapoor, 2017). For instance, mannanase from *B. subtilis* YH12 and B36 showed optimum activity at 55°C and 50°C, respectively (Liu et al., 2015). Notably, rGmuG retained over 80% of its enzyme activity at 60°C, indicating the potential for mannan hydrolysis to continue during the drying process of the fermented PKM at this high temperature.

In term of optimum pH, most bacterial β-mannanases was found maximally active at neutral to alkaline pH while fungal β-mannanases mostly show optimum activity at acidic pH. For instance, β-mannanase from *B. nealsonii* PN11 (GH5) and *Bacillus* N16-5 (GH26) showed optimum activity at pH 8 and 9.6, respectively (Srivastava & Kapoor, 2017). In contrast, β-mannanase derived from *B. subtilis* NM-39 and B36 worked optimally at pH 5.0 and 6.0, respectively. Interestingly, our rGmuG displayed two pH optima at acidic and alkaline pHs, respectively. This occurrence is reminiscent of the aspartic protease from *Penicillium roqueforti*, which exhibits maximal hydrolysis of casein at pH 3.5 and 5.5, possibly due to conformational changes in the substrate. In addition, the acidic lipase from *P. roqueforti* demonstrates a pH optimum at 6.0 and a less pronounced optimum at 2.8. (Cantor et al., 2004).

Lastly, the substrate specificity of rGmuG was examined using different synthetic or purified substrates. rGmuG was found to exhibit hydrolytic activity towards the different mannan substrates tested including 1,4-β-D-mannan, LBG galactomannan and konjac glucomannan. However, the enzyme displayed no activity with the various non-mannan substrates tested including xylan, sodium carboxymethyl cellulose and polygalacturonic acid. This further confirms our earlier hypothesis that rGmuG likely does not have cross-reactivity towards cellulose. Similarly, β-mannanase from *Penicillium oxalicum GZ-2* exhibited activity for LBG and guar gum galactomannan but not sodium carboxymethyl cellulose and beechwood xylan (Liao et al., 2014). In contrast, β-mannanase derived from *B. subtilis* YH12 exhibits an unusually broad substrate specificity (Liu et al., 2015). The enzyme not only degrades mannan, glucomannan, and galactomannan but also hydrolyses other polysaccharides with complex structures such as xanthan gum and carrageenan.

## 4. Conclusion

In this study, we isolated a high mannanase-producing strain of *B. subtilis* F6 from palm fruit exocarp through enrichment on near solid-state PKM over an extended duration, followed by screening based on mannanase production levels. The isolated strain was successfully applied in the fermentation of PKM, resulting in a substantial decrease of at least 10% in the NDF content after an overnight fermentation. Notably, mannanase activity was detected as early as 6 h into incubation, indicating a rapid response in mannanase production upon contact with PKM. Hence, the isolated *B. subtilis* strain F6 offers an effective solution for fibre degradation of PKM within a short fermentation time, reducing the risk of contamination and avoiding high production costs.

Subsequently, β-mannanase GmuG was identified as the primary carbohydrase responsible for the high mannanase activity and fibre hydrolysis capability of strain F6 during the fermentation process. The notable increase in extracellular mannanase activity was likely attributed to the concurrent upregulation of the glucomannan utilization operon and Sec translocase components. Furthermore, PKM hydrolyzed by rGmuG was enriched in mannobiose, followed by mannotriose and mannose. Hence, the fermentation process not only decreased the fibre content but also potentially enhanced the feed with valuable prebiotics and probiotics in the form of mannooligosaccharides and *B. subtilis*, respectively, transforming it into a functional feed ingredient.

With advancements in synthetic biology, there is potential to further enhance fibre degradation without requiring prolonged fermentation periods. Identification and characterization of mannanase GmuG involved in PKM fibre hydrolysis lay the foundation for future engineering of the bacterial strain. Moreover, additional insights can be gained by comparing the organoleptic properties and metabolites in F6-fermented PKM with those in PKM fermented using other microbes.

## Acknowledgements

The authors would like to thank Professor Chua Nam Hai for his constructive comments on this project, and Ms. Rachel Ng Swee Lin, Ms. Lee Pei Yin, and Ms. Caroline Tham Jun Ying for their technical assistance, and Mr. Hermil Calasang for arrangement of PKM sample.

## Funding

This work was supported by Wilmar International Ltd (WIL) as a research and development project. Wei Li Ong is supported by Singapore Economic Development Board Industrial Postgraduate Programme (EDB-IPP).

## Notes

### Competing Interest Statement

The authors have declared no competing interest.

